## Supplementary figures and images for "Dynamics of *Trypanosoma cruzi* infection in hamsters and novel association with progressive motor dysfunction"

### Nose_cone_back.png

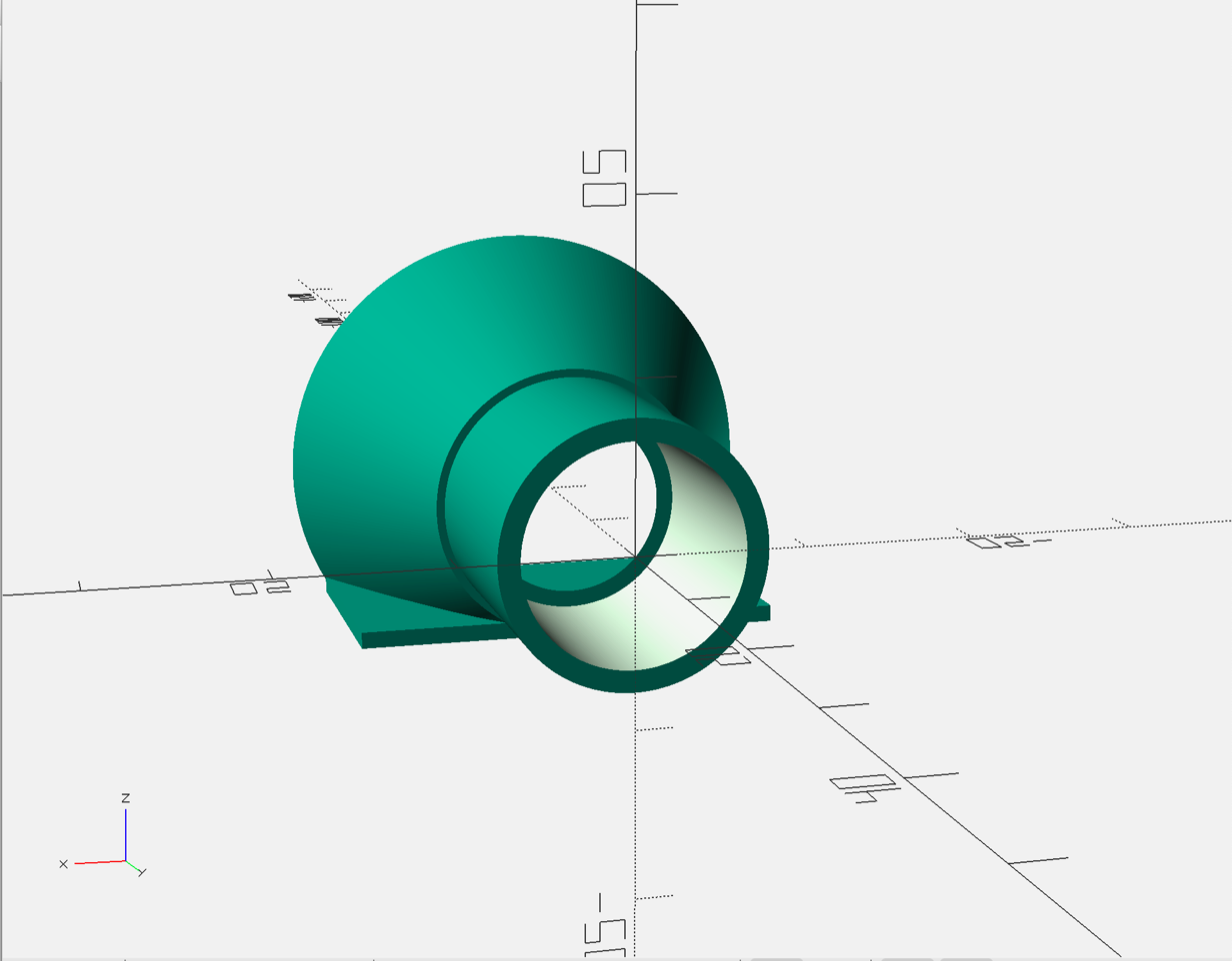

### Nose_cone_front.png

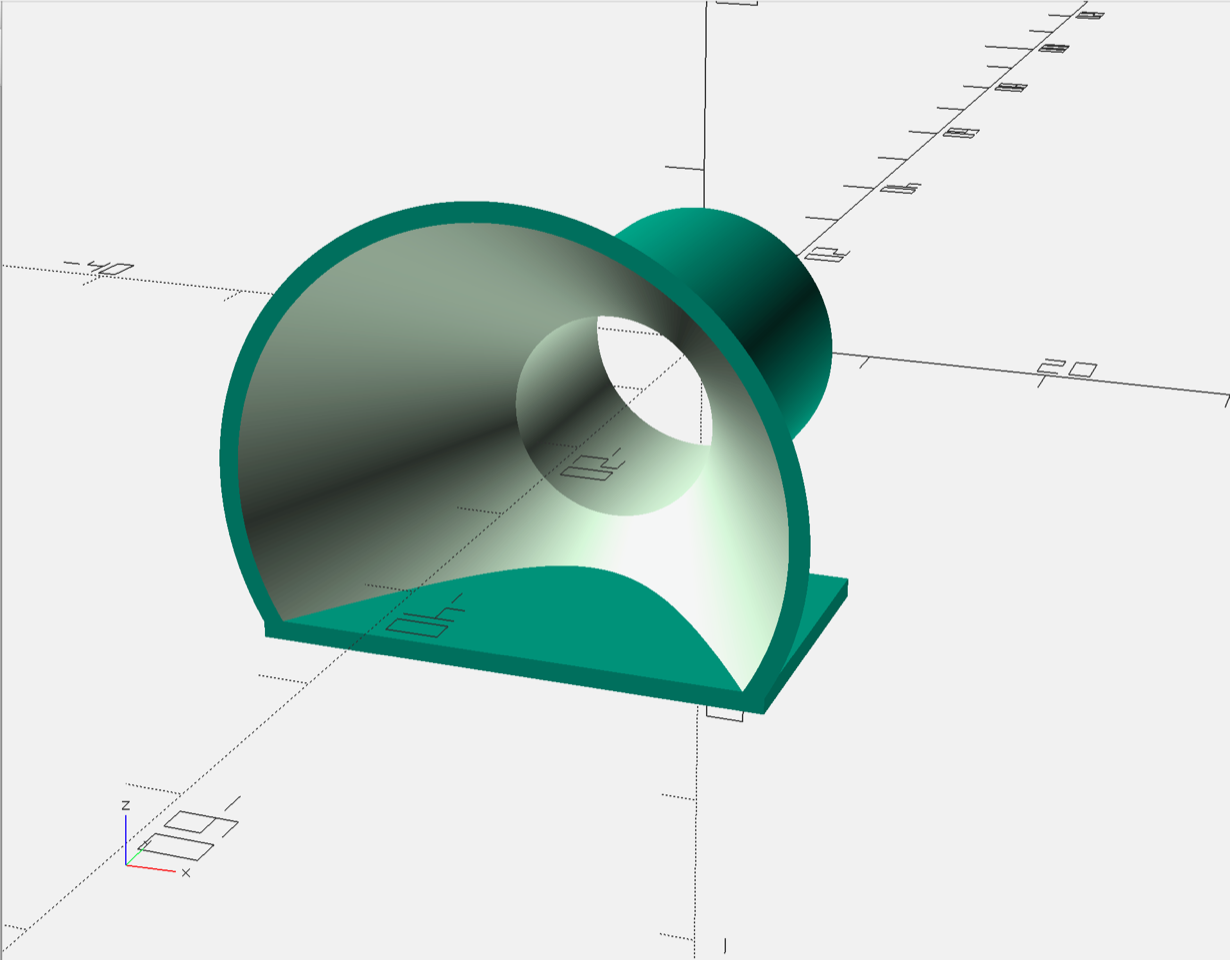
